# Detecting clonemates in multicellular clonal species based on “Shared Heterozygosity (SH)” and technical replicates

**DOI:** 10.1101/2022.02.16.480681

**Authors:** Lei Yu, John J. Stachowicz, Katie DuBois, Thorsten B. H. Reusch

## Abstract

Clonal reproduction, the formation of new individuals that are genetically nearly identical to the parent via mitosis in the absence of genetic recombination, is a very common reproductive mode across plants, fungi, and animals. Current genetic marker based methods fail to detect clonal structure when all collected samples belong to one single clone, which seems to be more common than previously thought. Here we propose a new similarity index, “Shared Heterozygosity (SH)” based on the number of genetic markers (typically SNPs, single-nucleotide polymorphisms) that are identically heterozygous among two or more genomes (i.e., N_SH_). Ideally N_SH_ should be on the order of approximately >=3,000, which can be easily achieved nowadays via Restriction-site Associated DNA (RAD) sequencing or whole-genome resequencing. One should be careful when N_SH_ is small (roughly <1,000). We analyze two large seagrass clones (*Posidonia australis, Zostera marina*) along with two *Z. marina* clones of known age (17-yrs), and show that SH can potentially extend the detection of clonemates to any pair of samples with the aid of technical replicates. Another potential application of SH is to detect possible parent-descendant pairs under selfing, because the heterozygous loci in the selfing-produced descendants represent a subset of those in the parent. Our proposed workflow takes advantage of the availability of the larger number of genetic markers in the genomic era, and fills a gap in detecting clonemates in the growing number of cases where many or all samples at a location belong to one single clone.

## Introduction

Clonal reproduction refers to the formation of new individuals that are genetically nearly identical to the parent via mitosis in the absence of genetic recombination (De Meeûs et al., 2007). This reproductive mode is widespread in multicellular organisms, and has many different forms, such as vegetative multiplication (animals and plants), parthenogenesis (animals), apomixis (plants), and gynogeneis or pseudogamy (De Meeûs et al., 2007; Orive & Krueger-Hadfield, 2021). In multicellular clonal species, a module or ramet (Harper & White, 1974) is the basic modular unit with an ability to live independently, while a genet (synonymous with clone) is the product of a single zygote from gamete fusion (Noble et al., 1979), and comprises all ramets that emerge via mitosis (i.e., clonemates) during its lifetime. The majority of multicellular clonal species can conduct both clonal and sexual reproduction (Schön et al., 2009), which is called partial clonality (Stoeckel et al., 2021a; Stoeckel et al., 2021b). Partially clonal taxa thus have two levels of biological organization, an asexual population of ramets belonging to a single clone/genet, which is nested into the population of clones/genets, the “classical”, sexually reproducing population (Reusch et al., 2021).

Ramets belonging to the same clone vs. different clones are mostly phenotypically indistinguishable and can also rarely be physically tracked once they obtain autonomy and break off the parental module. Hence, genetic markers such as microsatellites or AFLP have been widely used for detecting clonemates (Arnaud-Haond et al., 2007). A ramet can be genotyped with multiple independent markers, and the genotypes at all these loci collectively form a multilocus genotype (MLG). In case of co-dominant markers such as microsatellites, identical or nearly identical MLGs are then interpreted as being members of the same clone or genet. However, identical or nearly identical MLGs are not definitive evidence for clonal reproduction (Halkett et al., 2005), if for example there is low (or no) allelic diversity at particular loci. In order to calculate the probability of finding identical MLGs resulting from distinct zygotes, the population allelic frequencies are required, and the population is assumed to be under Hardy-Weinberg equilibrium (Parks & Werth, 1993). Evidently, estimates of such population-level frequencies are inaccurate in populations dominated by a few clones (Arnaud-Haond et al., 2007), and meaningless if a single site is dominated by one clone (Smith et al., 1992; Reusch et al., 1999; Arnaud-Haond et al., 2012; Japaud et al., 2015; Edgeloe et al., 2022), thereby rendering distinction of clonemates from non-clonemates difficult.

Another difficulty arises because members of the same clone may have slightly different MLGs (Douhovnikoff & Dodd, 2003; Yu et al., 2020; Edgeloe et al., 2022) owing to somatic genetic polymorphisms or technical errors. To overcome this issue and also to work with dominant markers that do not allow for the application of MLGs, an alternative suite of approaches is the analysis of pairwise genetic distances/similarities. The genomes of the ramets from the same clone are identical by descent (IBD), except at those sites that have experienced somatic mutations, as the genomes are inherited from the same asexual ancestor via mitosis without recombination. Thus, ramets from the same clone will show much lower levels of genetic distances or much higher levels of genetic similarities than those from different clones. This will lead to a bimodal pattern in the frequency distribution of pairwise genetic distances/similarities. Then, a threshold can be defined to separate the intra-clone vs. inter-clone ramet pairs (Douhovnikoff & Dodd, 2003; Bailleul et al., 2016; Meirmans, 2020). Such a threshold can be defined arbitrarily by the user, or by simulating a pseudo-observed generation of pure sexual reproduction from the genotyped population (Bailleul et al., 2016). Obviously, one problem is that the collected samples have to belong to more than one clones/genets. Otherwise, it would be impossible to identify the intra-clone sample pairs.

Here, we propose a novel genetic similarity index, Shared Heterozygosity (SH), which corresponds to the number of heterozygous sites shared by all individuals (ramets) divided by the number of heterozygous sites observed in the individual with the highest count. SH is an extension to Jaccard similarity index, with some modifications. SH involves a selection of marker loci, taking into account only the ones where at least one sample is heterozygous. Homozygous sites are usually not informative, as even distantly related individuals are identically homozygous at most of the genomic sites (The 1000 Genomes Project Consortium, 2015; The 1001 Genomes Consortium, 2016). On the contrary, heterozygous sites are more informative for detecting clonemates, because they show the potential for segregation in sexual reproduction, while remaining identically heterozygous in clonal reproduction.

The genetic differences between clonemates are determined by both technical errors and somatic genetic polymorphisms. By analyzing some of the largest detected clones ever (Yu et al., 2020; Edgeloe et al., 2022), we provided a conservative estimate as to how large the within-clone divergence in terms of non-shared heterozygosity may get. Even for these very large (and possibly old) clones, the level of non-shared heterozygosity caused by somatic genetic polymorphisms was low. Therefore, after having quantified the influence of technical errors with the aid of technical replicates (i.e., multiple pieces of tissues from the same sample are used for all the processes from DNA extraction to genotype calling), it may be possible to identify the intra-clone sample pairs without inter-clone information, which will extend the clonemate detection to any pair of samples.

Our workflow is most suitable for large number of SNPs obtained from whole-genome resequencing, restriction-site associated DNA (RAD) sequencing, or SNP arrays. The application of SH requires a sufficient number of genetic markers where the two target samples are identically heterozygous (i.e., N_SH_, ideally on the order of >= 3,000). Since the number of markers used does not always reflect the level of N_SH_, one should pay attention to N_SH_ and be careful when the level of N_SH_ is low (roughly <1,000). The goal of this study is to develop the workflow of detecting clonemates based on shared heterozygosity and technical replicates (Fig. 1). We apply our workflow to two of the largest clones in marine systems detected so far, seagrass clones in Finland (*Zostera marina*, covering >300 m x 200 m) and Australia (*Posidonia australis*, spanning >180 km), respectively (Yu et al., 2020; Edgeloe et al., 2022), which likely represent the far end of accumulating somatically generated variation. We also apply the workflow to two 17-year-old *Z. marina* clones cultured in outdoor flow-through mesocosms, representing the near end of accumulating somatically generated variation.

**Figure 1:**
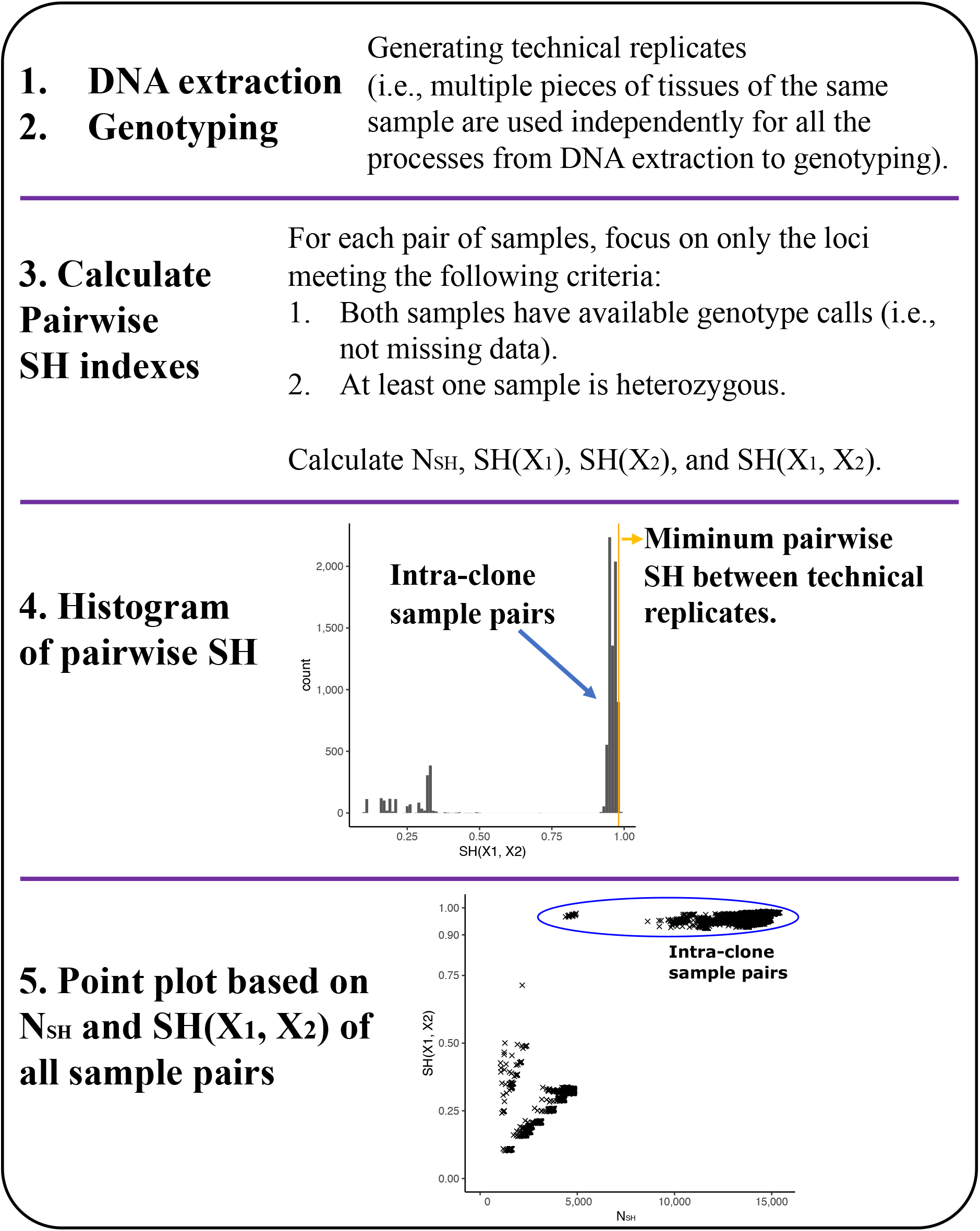
Workflow for detecting clonemates in multicellular clonal species based on SH indexes and technical replicates.

## Materials and Methods

### Shared Heterozygosity (SH) indexes

For n samples in multicellular clonal species (X_1_, …, X_n_), N_SH_(X_1_, …, X_n_) denotes the number of the genetic markers where n samples are identically heterozygous; N_Het_(X_n_) denotes the number of the heterozygous loci in X_n_.

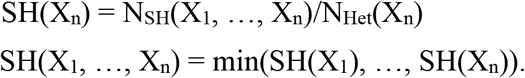

### Detecting clonemates based on a published dataset on seagrass *Posidonia australis*

Edgeloe et al. (2022) reported a polyploid *P. australis* clone spanning at least 180 km. The SNP dataset (all.vcf) containing 18,021 SNPs for 133 individuals (including 4 technical replicates) from 10 populations was used here. According to Edgeloe et al. (2022), population GU was diploid, while the other 9 populations were polyploid and belonged to the same clone. SH indexes were calculated for each pair of samples based on the SNP dataset, and clonemate detection was conducted following the workflow illustrated by Fig. 1.

### Distribution of SH indexes within two *Z. marina* clones that are 17 years old

Two *Z. marina* clones were selected from the eight clones cultured in adjacent 300-l outdoor flow-through mesocosms at Bodega Marine Laboratory (BML) under ambient light and temperature conditions (Hughes et al., 2009). Each mesocosm was initiated with a single terminal shoot collected from one of eight different locations in Bodega Harbor, California, in July 2004. The genetic distinctiveness of each of the original shoots was verified before planting. Six shoots (= ramets) were collected from each clone for genomic analysis in 2021, after the clones had grown in mesocosm for 17 years. The two clones were referred to as “GREEN” and “RED”, respectively.

Genomic DNA was extracted using NucleoSpin plant II kit from Macherey-Nagel following the instructions. The plant tissue (160-200 mg fresh weight) was ground with liquid nitrogen. Library construction and Illumina sequencing (PE150, paired end 150 bp) were conducted by BGI (Beijing Genomics Institute, Hong Kong). The quality of the raw Illumina reads was assessed by FastQC V0.11.5 (https://www.bioinformatics.babraham.ac.uk/projects/fastqc/). BBDuk (last modified November 7, 2019, https://jgi.doe.gov/data-and-tools/bbtools/bb-tools-user-guide/bbduk-guide/) was used to remove adapter contamination and conduct a basic quality filtering: (1) reads with more than one N were discarded (maxns=1); (2) reads shorter than 50 bp after trimming were discarded (minlength=50); (3) reads with average quality below 10 after trimming were discarded (maq=10). FastQC V0.11.5 was used for second round of quality check for the filtered reads.

The quality-filtered Illumina reads were mapped against the chromosome-level *Z. marina* reference genome V3.1 (Ma et al., 2021) using BWA MEM V0.7.17 (Li & Durbin, 2009). The alignments were converted to BAM format and then sorted using Samtools V1.11 (Li et al., 2009). The MarkDuplicates module in GATK V4.1.1.0 (Van der Auwera & O’Connor, 2020) was used to identify and tag duplicate reads in the BAM files. Sequencing depth and percentage of covered genome region for each ramet were calculated using Samtools V1.11 and Bedtools V1.11 (Quinlan & Hall, 2010), respectively. HaplotypeCaller (GATK V4.1.1.0) was used to generate GVCF format file for each ramet, and all the GVCF files were combined by CombineGVCFs (GATK V4.1.1.0). GenotypeGVCFs (GATK V4.1.1.0) was used to call genetic variants.

BCFtools V1.11 (Li, 2011) was used to remove SNPs within 20 base pairs of an indel or other variant type. Then we extracted only bi-allelic SNPs located in the six main chromosomes (2,969,820 SNPs). INFO annotations were extracted using VariantsToTable (GATK V4.1.1.0). SNPs meeting one or more than one of the following criteria were marked by VariantFiltration (GATK V4.1.1.0): MQ < 40.0; FS > 60.0; QD < 13.0; MQRandSum > 2.5 or MQRandSum < −2.5; ReadPosRandSum < −2.5; ReadPosRandSum > 2.5; SOR > 3.0; DP > 1192.45 (2 * average DP). Those SNPs were excluded by SelectVariants (GATK V4.1.1.0). A total of 1,105,628 SNPs were retained. VCFtools V0.1.15 (Danecek et al., 2011) was used to convert individual genotypes to missing data when GQ < 30 or DP < 20. Individual homozygous reference calls with one or more than one reads supporting the variant allele, and individual homozygous variant calls with one or more than one reads supporting the reference allele, were also converted to missing data using a custom Python script (https://github.com/leiyu37/Detecting-clonemates.git). Finally, we kept only genomic sites showing polymorphism (604,066 SNPs).

SH indexes were calculated by a custom Phython script (https://github.com/leiyu37/Detecting-clonemates.git) for each pair of samples. We only focused on the nucleotide positions in the reference genome where both samples had available genotype calls (not missing data).

### Detecting clonemates based on a published dataset on seagrass *Zostera marina*

Yu et al. (2020) conducted whole-genome resequencing of 24 ramets of a seagrass *Z. marina* clone located in Finland. The clone was estimated to be between 750 and 1,500 years old. Here the clone was referred to as “the Finnish clone”. All the 24 ramets were sequenced by Illumina Hiseq, and three of them were also sequenced independently by Illumina Novaseq. The Hiseq and Novaseq data for the three ramets provided three pairs of technical replicates. The Novaseq data reached a coverage of ∼1000x. A subset was selected from the Novaseq data (∼80x) to achieve a similar coverage to the Hiseq data. Then we re-analyzed the Hiseq and Novaseq (∼80x) data together using the same method above for the two 17-year-old clones. All the SNP calling and filtering processes were exactly same with those for the 17-year-old clones, except the threshold for DP during hard filtering (DP > 2945.57). SH indexes were calculated for each pair of samples.

To understand the influence of N_SH_(X_1_, X_2_) on SH(X_1_, X_2_), we randomly selected 1/10, 1/100 and 1/1000 from the 38,831 SNPs for the 24 ramets available in Yu et al. (2020), based on which N_SH_(X_1_, X_2_) and SH(X_1_, X_2_) were calculated again for each pair of samples. Each selection was repeated 100 times, and N_SH_(X_1_, X_2_) and SH(X_1_, X_2_) from the 100 repeats were pooled together to analyze.

## Results

We applied our workflow to two of the largest clones in marine systems detected so far, seagrass clones in Finland (*Zostera marina*, covering >300 m x 200 m) and Australia (*Posidonia australis*, spanning >180 km), respectively (Yu et al., 2020; Edgeloe et al., 2022). In the larger clone of *P. australis* in Shark Bay (Western Australia), SNPs were obtained by RAD (Edgeloe et al., 2022). Based on the SNP dataset encompassing 18,021 SNPs for 133 individuals (including 4 technical replicates) from 10 populations, the pairwise N_SH_ ranged from 1,047 to 15,428. The minimum pairwise SH between technical replicates was 0.9809. The pairwise SH between clonemates were smaller than 0.9809, due to somatic genetic variation. They were distributed within a narrow range bordered by the technical replicates (Fig. 2a). Including technical replicates made it possible to identify intra-clone sample pairs without inter-clone information (Fig. 2a). Our analyses based on SH indexes and technical replicates confirmed that all populations except population GU belonged to the same clone (Fig. 2b).

**Figure 2:**
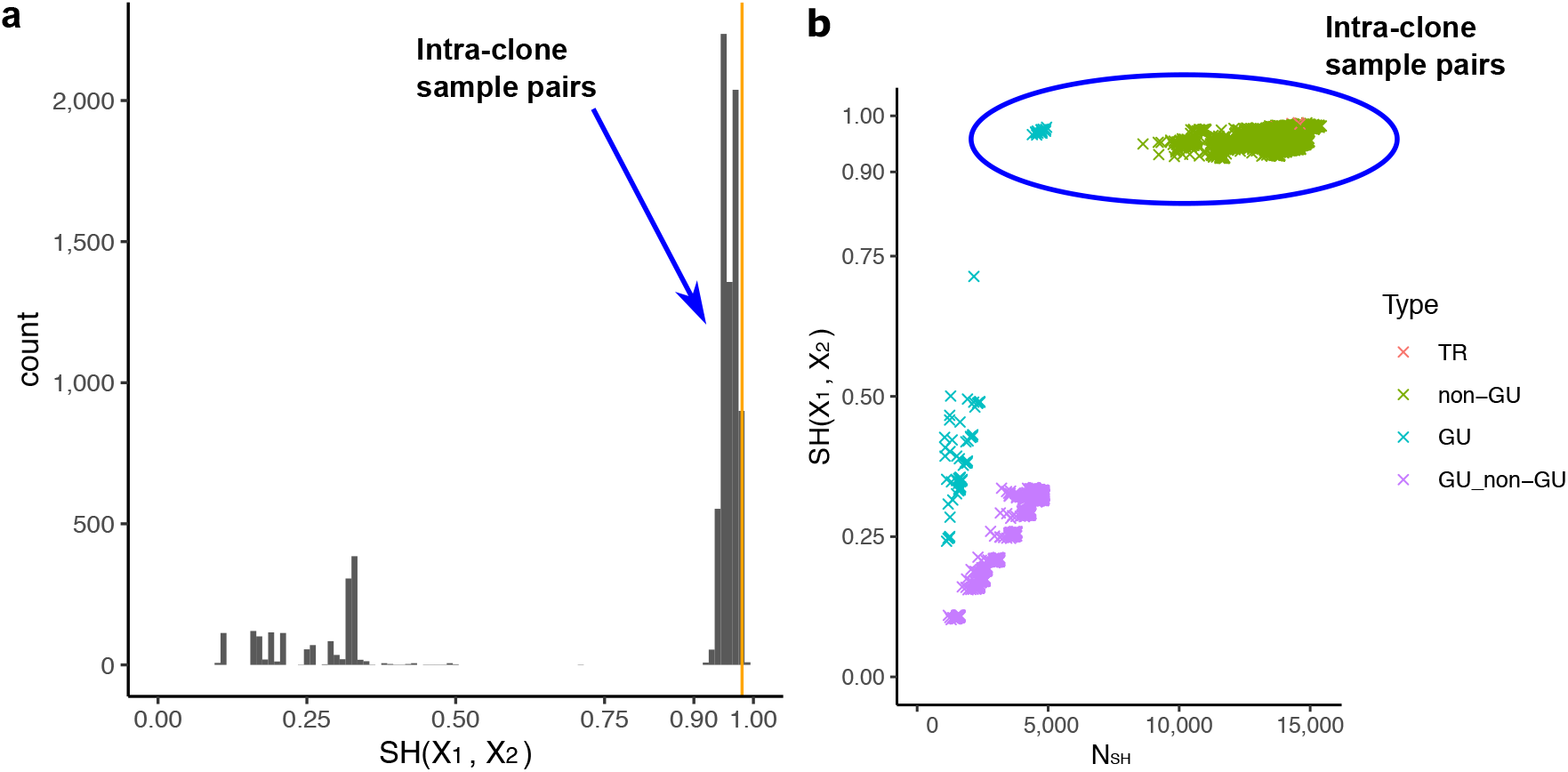
Detecting clonemates using SH indexes and technical replicates based on a published dataset on seagrass *Posidonia australis*. **a, Histogram of pairwise SH**. The dataset contains four pairs of technical replicates. The orange vertical line indicates the minimum pairwise SH between technical replicates. The pairwise SH between clonemates are expected to be smaller than the minimum pairwise SH between technical replicates, due to somatic genetic variation. They form a distribution to the left of the orange vertical line. **b, Point plot based on all sample pairs**. Each data point represents one pair of samples. The sample pairs are categorized into four different types: 1, TR: a pair of technical replicates for the same sample; 2, non-GU: neither samples are from population GU; 3, GU: both samples are from population GU; 4, GU_non-GU: one sample is from population GU, and the other sample is from the other 9 populations.

In the relatively smaller clone of *Z. marina* in Finland, whole-genome resequencing was conducted for 24 ramets (Yu et al., 2020). After re-analyzing the data (Hiseq data for 24 ramets, and Novaseq data for 3 of the ramets), we obtained 41,885 SNPs. The Hiseq and Novaseq data for the three ramets provided three pairs of technical replicates. Based on the dataset, the pairwise N_SH_ ranged from 28,619 to 32,210 (Fig. 3). The minimum pairwise SH between technical replicates was 0.9998 (Fig. 3a). The pairwise SH between clonemates were distributed within a narrow range from 0.9548 to 0.9993, bordered by the technical replicates (Fig. 3b). Our analyses based on SH indexes and technical replicates confirmed that all 24 ramets belonged to the same clone. By calculating SH indexes based on the the subsets of the SNP dataset, we found that smaller N_SH_ led to larger variation of pairwise SH (Fig. 4). The consistently high SH when N_SH_ is at least on the order of >=3,000 suggests this as a reasonable minimum target to apply this method. When N_SH_ <1,000, variability in SH increased considerably, which would reduce confidence in distinguishing clonemates from non-clonemates.

**Figure 3:**
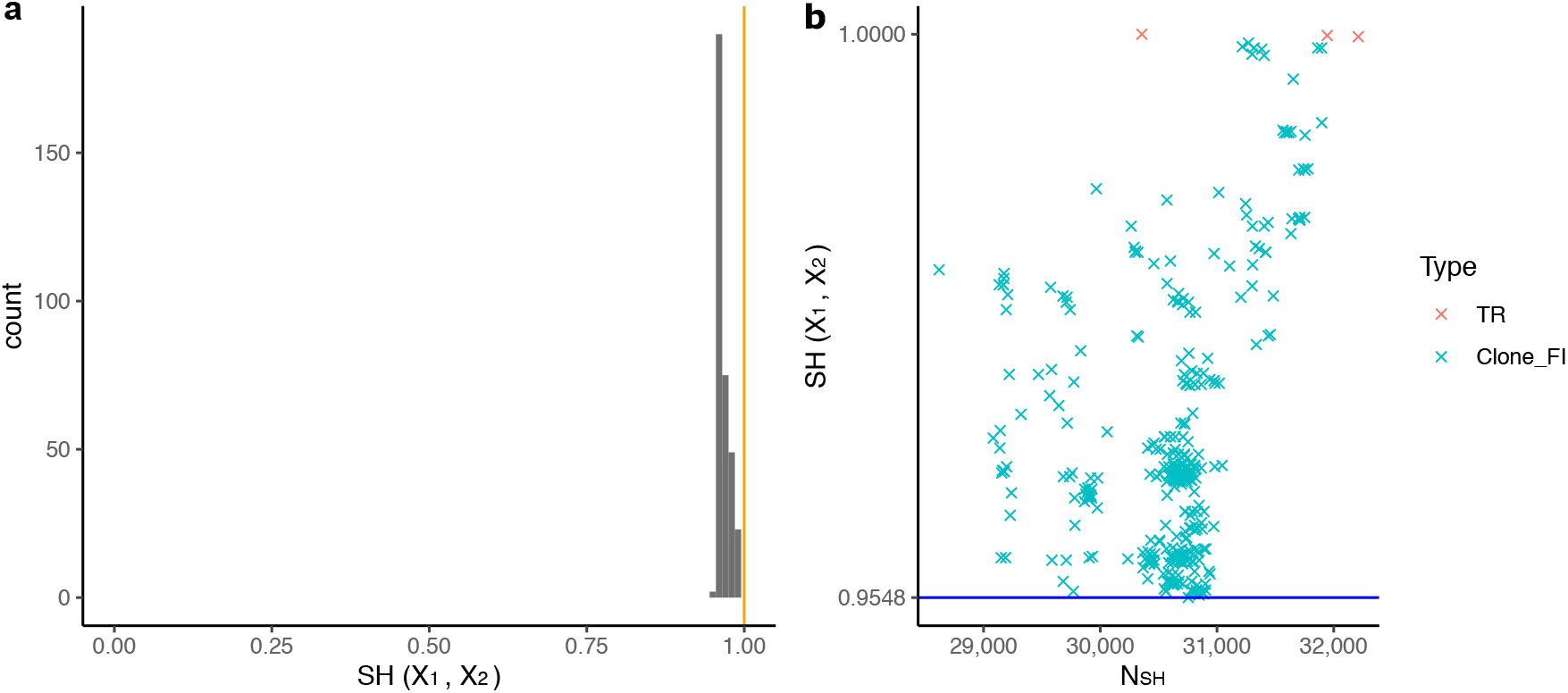
Detecting clonemates using SH indexes and technical replicates based on a published dataset on seagrass *Zostera marina*. **a, Histogram of pairwise SH**. The dataset contains 24 ramets of the same clone. Three of them were sequenced independently on both Illumina Hiseq and Novaseq platform, providing three pairs of technical replicates. The orange vertical line indicates the minimum pairwise SH between technical replicates. The pairwise SH between clonemates are expected to be smaller than the minimum pairwise SH between technical replicates, due to somatic genetic variation. They form a distribution to the left of the orange vertical line. **b, Point plot based on all sample pairs**. Each data point represents one pair of samples. The sample pairs are categorized into two different types: 1, TR: a pair of technical replicates for the same sample; 2, Clone_FI: the samples pairs that do not belong to TR. The blue line indicates the minimum pairwise SH within Clone_FI.

**Figure 4:**
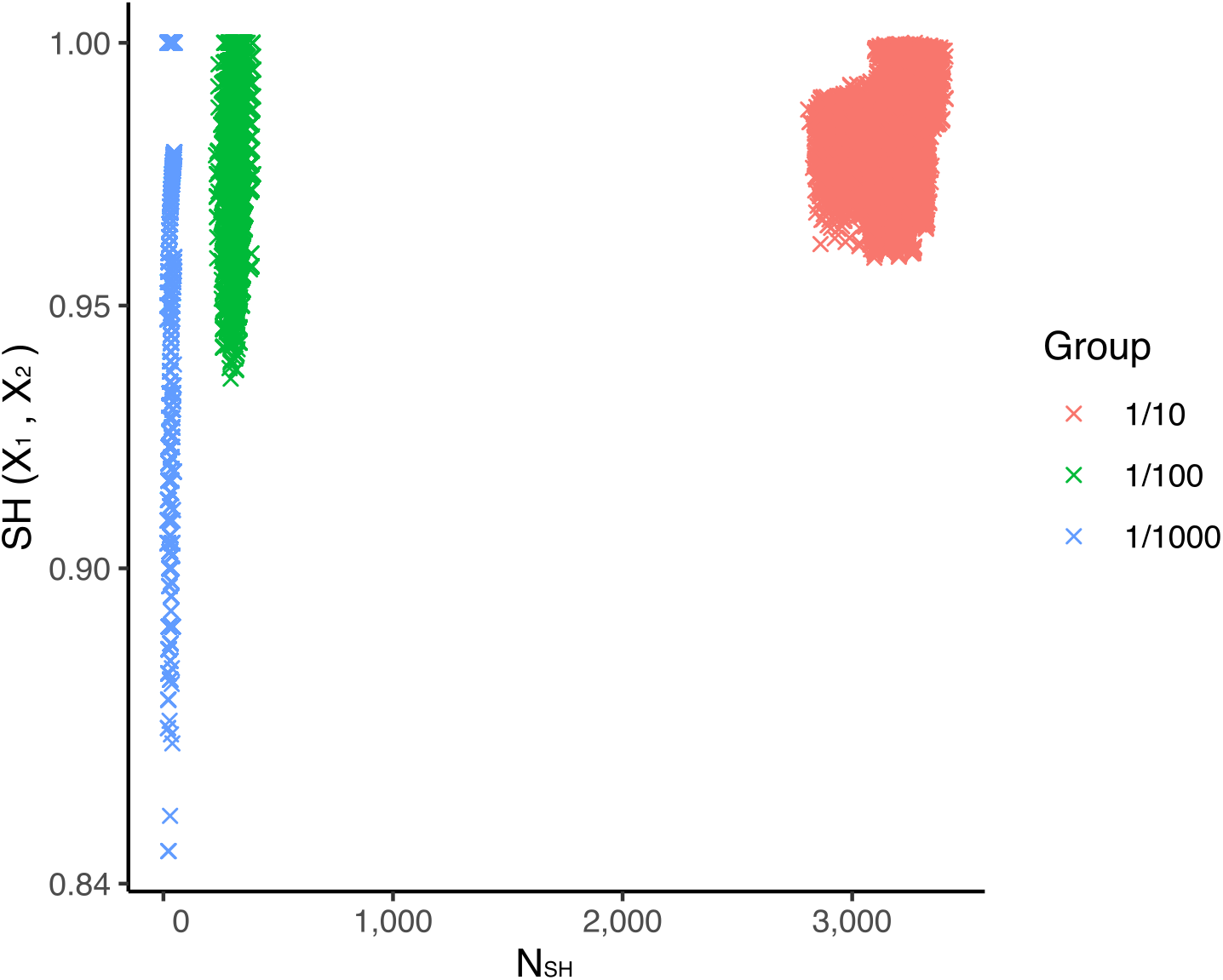
Distribution of pairwise SH indexes within the Finnish *Zostera marina* clone based on subsets (1/10, 1/100, and 1/1000) of SNPs. The variation of SH(X_1_, X_2_) increases when N_SH_ decreases.

We also calibrated our approach using 2 relatively young *Z. marina* clones of known age (17-year-old clones). Unfortunately, technical replicates were also not available. The whole-genome resequencing data of the 12 cultured *Z. marina* ramets covered >93% of the reference genome, and reached >66x coverage (Table 1). The pairwise N_SH_ ranged from 108,585 to 288,606 (Fig. 5a). Although technical replicates were not available, the pairwise SH between technical replicates should be distributed within the range from 0.9980 to 1 (Fig. 5b). Within the clone “GREEN”, the minimum SH(X_1_, X_2_) was 0.9975 (Fig. 5a). As for the clone “RED”, we detected one “contaminant” ramet (i.e., R10), obviously not belonging to that clone. The five ramet pairs composed of R10 and one of the other five “RED” ramets showed SH(R10, X_2_) ranging from 0.4103 to 0.4114 (Fig. 5a), indicating that R10 does not belong to this clone. Apart from R10, the minimum SH(X_1_, X_2_) for the ramet pairs among the other five “RED” ramets was 0.9974 (Fig. 5a). The inter-clone SH(X_1_, X_2_) ranged from 0.4790 to 0.4803. Since the pairwise SH between clonemates were distributed within a narrow range from 0.9974 to 0.9980 (Fig. 5b), they could be easily identified.

**Table 1:** Coverage of the whole-genome resequencing data for the two cultured *Zostera marina* clones.

| Sample | Average depth for the covered region | Genome region covered (%) |
| --- | --- | --- |
| G01 | 68.64 | 93.33 |
| G02 | 80.42 | 93.66 |
| G06 | 79.05 | 93.72 |
| G07 | 69.94 | 93.36 |
| G10 | 77.49 | 93.52 |
| G16 | 73.31 | 93.26 |
| R02 | 81.55 | 93.86 |
| R03 | 85.13 | 94.06 |
| R05 | 68.14 | 93.57 |
| R08 | 66.44 | 93.51 |
| R09 | 71.73 | 93.59 |
| R10 | 67.97 | 93.69 |

**Figure 5:**
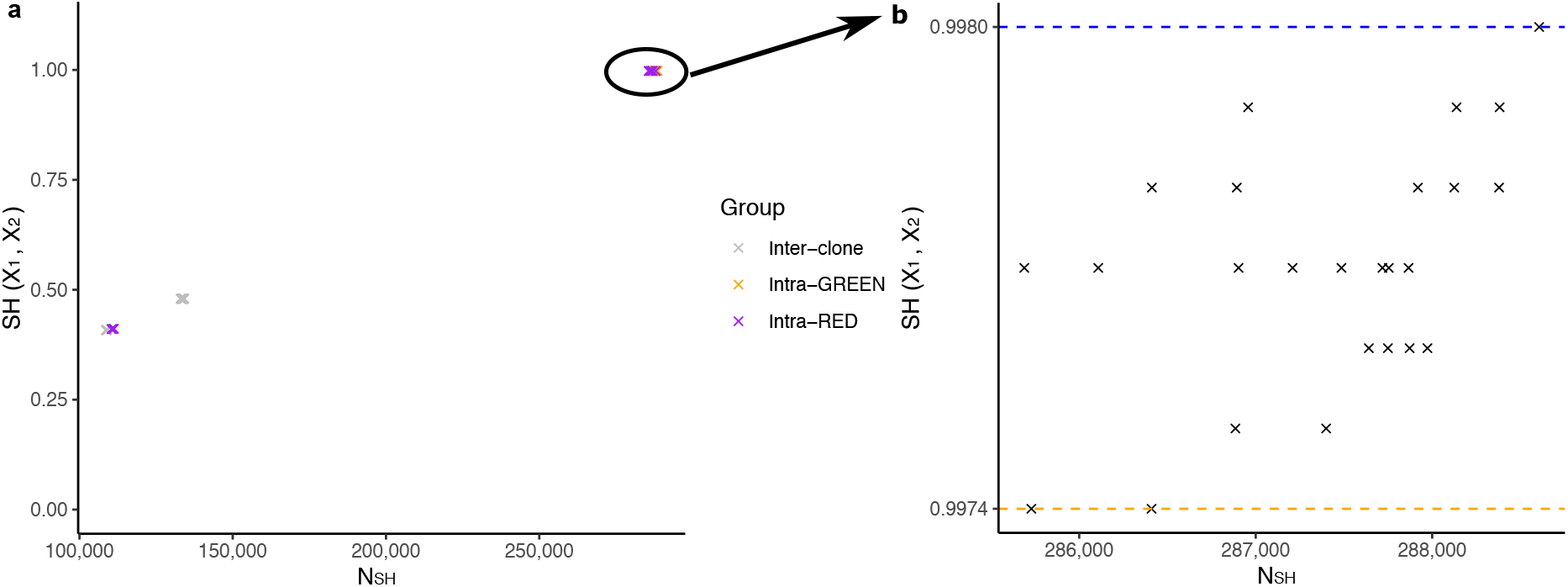
Distribution of pairwise SH indexes within the two 17-year-old *Zostera marina* clones. **a, SH indexes for all sample pairs**. Each data point represents one sample pair, which is either from the same clone (“intra-GREEN” or “intra-RED”) or from different clones (“inter-clone”). One ramet from clone “RED” (i.e., R10) obviously belongs to another clone, leading to much lower levels of pairwise SH with ramets of clone “RED”. **b, SH indexes for sample pairs of the same clone**. The orange and the blue dotted lines indicate the minimum and maximum values for SH(X_1_, X_2_), respectively.

## Discussion

Both sexual and clonal reproduction occur in almost all clonal taxa, but the length and extent of the clonal phase varies widely within and among species (for seagrasses, e.g. Reusch & Boström, 2011). Examples accumulate that some populations can propagate clonally to such an extent that the whole population is dominated by one or a few clones, such as fungi (Smith et al., 1992), seagrass species *Zostera marina* (Reusch et al., 1999; Yu et al., 2020), *Posidonia oceanica* (Arnaud-Haond et al., 2012) and *P. australis* (Edgeloe et al., 2022), and corals (Japaud et al., 2015). The situation of having a collection of samples belonging to one single clone (Yu et al., 2020) is thus more widespread than previously thought. The currently available methods cannot work in this extreme case (Arnaud-Haond et al., 2007). We have proposed a workflow based on the concept of “Shared Heterozygosity (SH)”, meaning that two or more than two genomes are identically heterozygous at some genetic markers. The pairwise SH between clonemates are determined by both technical errors and somatic genetic variation. However, even for the two very old seagrass clones, the level of pairwise differences caused by somatic genetic variation is relatively low (Fig. 2&3). We have shown that intra-clone sample pairs can be easily identified with the aid of the technical replicates even without inter-clone information (Fig. 3). Our workflow can be applied to any pair of ramets when technical replicates are available, even if there is only one genet occurring within a “population”-level sampling. To the best of our knowledge, this is the only method that can work in cases where all collected ramets belong to one single clone.

When applying the workflow to the two 17-year-old *Z. marina* clones, we found one “contaminant” ramet (i.e., R10) that clearly did not belong to the clone “RED”. One possible reason is that R10 was produced by sexual event between members of the clone “RED” (i.e., selfing). In the first 15 years of growth, this would be the only way for contamination to occur, as the clones were cultured in isolated containers. Under selfing, each of the heterozygous loci in the parent can either remain identically heterozygous or change to a homozygous state in R10. As a consequence, the heterozygous loci in R10 should be a subset of those in the parent, and thus SH(R10) for the five ramet pairs composed of R10 and one of the other five “RED” ramets should be close to 1. However, that is not the case, as SH(R10) for the five ramet pairs ranges from 0.4320 to 0.4331. Hence, R10 is not likely to be derived from selfing. In late 2019 or early 2020, these clones were transferred and planted in separate containers within in a single larger tank (∼4 m diameter). All clones were in the same tank for at most one year. It is possible that either (a) outcrossing occurred and seeds set and germinated or (b) shoots dislodged and re-attached or were able to grow between pots.

SH can potentially be used to detect possible parent-descendant pairs under selfing, because this reproductive mode leads to loss of heterozygosity. Under selfing, a subset of the heterozygous loci in the parent will change to a homozygous state in the descendants. If not considering the ramet-specific heterozygous loci derived from mutations and genotyping errors, the heterozygous loci in the descendant should be a subset of those in the parent, which can potentially be used to detect possible parent-descendant pairs under selfing.

The application of SH requires a sufficient number of genetic markers where the two target samples are identically heterozygous (i.e., N_SH_). We were able to correctly detect clonemates based on the datasets with minimum N_SH_ of 1,047. With the development of sequencing technology, it is now easy to obtain thousands of genetic markers using whole-genome resequencing, restriction-site associated DNA (RAD) sequencing, or SNP arrays. However, the level of N_SH_ can still be low based on genome-wide maker sets, when the target species shows very low levels of genome heterozygosity. One should pay attention to N_SH_ and use caution in the interpretation of results when N_SH_ is low. Our workflow takes advantage of the availability of the larger number of genetic markers in the genomic era. It can potentially extend the detection of clonemates to any pair of samples against the baseline of technical replicates and fills a gap in detecting clonemates in extreme cases where all sampled genotypes at a site belong to one single clone.

## Acknowledgements

This study was supported by a 4-year PhD scholarship from the China Scholarship Council (CSC) to L.Y. (No. 201704910807), and by a Human Frontiers of Science (HFSP) grant to T.B.H.R (RGP0042_2020). We thank D. Gill for lab assistance. We thank Dr. Y. Li for the comments on the early version of the manuscript.

## Data Accessibility

DNA sequence data are available in the NCBI short read archive, BioProject no. PRJNA806459, SRA accession no. SRR18000159-SRR18000170. Custom-made computer code is available at Github https://github.com/leiyu37/Detecting-clonemates.

## Authors’ contributions

L.Y. and T.B.H.R. conceived and designed the study, J.J.S and K.D. cultured and provided the samples of the two 17-year-old seagrass *Zostera marina* clones, L.Y. extracted DNA and analyzed the sequencing data, L.Y. performed the bioinformatic analyses, L.Y. and T.B.H.R. wrote the paper, with input from all authors.

## Samping permit

The original sampling of *Zostera marina* clones was conducted under the permit no. SC-4874.

## Competing interests

The authors declare that they have no competing interests.

## Notes

### Competing Interest Statement

The authors have declared no competing interest.

### Summary of Updates

1. The term identity by heterozygosity (IBH) is confusing, so we have decided to change it to shared heterozygosity (SH). 2. Giving an arbitrary threshold of 0.95 does not make sense, so we now use technical replicates to help identify intra-clone sample pairs. 3. We also try to give a clearer definition of shared heterozygosity (SH). 4. We re-analyzed the raw Next-Generation sequencing data in Yu et al. (2020), instead of directly using the available vcf file. This was because the vcf file was based on the Illumina Hiseq data for the 24 ramets. There were also Illumina Novaseq data available for 3 of the 24 ramets, which were used for detecting mosaic somatic genetic variation in Yu et al. (2020). The Hiseq data and Novaseq data for the three ramets provided three pairs of technical replicates, which was important for our proposed method. Hence, we re-analyzed all those data. 5. We realized that our method is also appliable to polyploid clones, so we have removed 'diploid' from the manuscript. We have added the analyses on a polyploid clone of another species. 6. We try to discuss/credit previous work relevant to our study, such as IBD (Identity by Descent) and Jaccard index. 7. We have replaced previous Figure 1 with a workflow for our method, added a new figure (Fig. 2) for the analyses on the newly added species, and modified the visualization for the other figures.

## References

Arnaud-Haond, S., Duarte, C. M., Diaz-Almela, E., Marbà, N., Sintes, T. & Serrão, E. A. (2012). Implications of extreme life span in clonal organisms: millenary clones in meadows of the threatened seagrass Posidonia oceanica. PLoS ONE, 7(2), e30454. doi:10.1371/journal.pone.0030454

Arnaud-Haond, S., Duarte, C. M., Alberto, F. & Serrao, E. A. (2007). Standardizing methods to address clonality in population studies. Mol. Ecol., 16(24), 5115–5139. doi:10.1111/j.1365-294X.2007.03535.x

Bailleul, D., Stoeckel, S. & Arnaud-Haond, S. (2016). RClone: a package to identify MultiLocus Clonal Lineages and handle clonal data sets in r. Methods Ecol. Evol., 7(8), 966–970. doi:10.1111/2041-210X.12550

Buss, L. W. (1983). Evolution, development, and the units of selection. Proc. Natl. Acad. Sci. U.S.A, 80(5), 1387–1391. doi:10.1073/pnas.80.5.1387

Danecek, P., Auton, A., Abecasis, G., Albers, C. A., Banks, E., DePristo, M. A., … Genomes Project Analysis Group. (2011). The variant call format and VCFtools. Bioinformatics, 27(15), 2156–2158. doi:10.1093/bioinformatics/btr330

De Meeûs, T., Prugnolle, F. & Agnew, P. (2007). Asexual reproduction: genetics and evolutionary aspects. Cell. Mol. Life Sci., 64(11), 1355–1372. doi:10.1007/s00018-007-6515-2

Douhovnikoff, V. & Dodd, R. S. (2003). Intra-clonal variation and a similarity threshold for identification of clones: application to Salix exigua using AFLP molecular markers. Theor. Appl. Genet., 106(7), 1307–1315. doi:10.1007/s00122-003-1200-9

Edgeloe, J. M., Severn-Ellis, A. A., Bayer, P. E., Mehravi, S., Breed, M. F., Krauss, S. L., … Sinclair, E. A. (2022). Extensive polyploid clonality was a successful strategy for seagrass to expand into a newly submerged environment. Proc. Royal Soc. B, 289(1976), 20220538. doi:10.1098/rspb.2022.0538

Halkett, F., Simon, J. & Balloux, F. (2005). Tackling the population genetics of clonal and partially clonal organisms. Trends Ecol. Evol., 20(4), 194–201. doi:10.1016/j.tree.2005.01.001

Harper, J. L. & White, J. (1974). The demography of plants. Annu. Rev. Ecol. Syst., 5(1), 419–463. doi:10.1146/annurev.es.05.110174.002223

Honnay, O. & Bossuyt, B. (2005). Prolonged clonal growth: escape route or route to extinction? Oikos, 108(2), 427–432. doi:10.1111/j.0030-1299.2005.13569.x

Hughes, R. A., Stachowicz, J. J. & Williams, S. L. (2009). Morphological and physiological variation among seagrass (Zostera marina) genotypes. Oecologia, 159(4), 725–733. doi:10.1007/s00442-008-1251-3

Japaud, A., Bouchon, C., Manceau, J. & Fauvelot, C. (2015). High clonality in Acropora palmata and Acropora cervicornis populations of Guadeloupe, French Lesser Antilles. Mar. Freshw. Res., 66(9), 847–851. doi:10.1071/MF14181

Li, H. (2011). A statistical framework for SNP calling, mutation discovery, association mapping and population genetical parameter estimation from sequencing data. Bioinformatics, 27(21), 2987–2993. doi:10.1093/bioinformatics/btr509

Li, H. & Durbin, R. (2009). Fast and accurate short read alignment with Burrows-Wheeler transform. Bioinformatics, 25(14), 1754–1760. doi:10.1093/bioinformatics/btp324

Li, H., Handsaker, B., Wysoker, A., Fennell, T., Ruan, J., Homer, N., … Genome Project Data Processing, S. (2009). The Sequence Alignment/Map format and SAMtools. Bioinformatics, 25(16), 2078–2079. doi:10.1093/bioinformatics/btp352

Ma, X., Olsen, J. L., Reusch, T. B. H., Procaccini, G., Kudrna, D., Williams, M., … Van de Peer, Y. (2021). Improved chromosome-level genome assembly and annotation of the seagrass, Zostera marina (eelgrass). F1000research, 10. doi:10.12688/f1000research.38156.1

Meirmans, P. G. (2020). genodive version 3.0: Easy-to-use software for the analysis of genetic data of diploids and polyploids. Mol. Ecol. Resour., 20(4), 1126–1131. doi:10.1111/1755-0998.13145

Noble, J. C., Bell, A. D. & Harper, J. L. (1979). The population biology of plants with clonal growth: I. The morphology and structural demography of Carex arenaria. J. Ecol., 67, 983–1008. doi:10.2307/2259224

Orive, M. E. & Krueger-Hadfield, S. A. (2021). Sex and Asex: A clonal lexicon. J. Hered., 112(1), 1–8. doi:10.1093/jhered/esaa058

Parks, J. C. & Werth, C. R. (1993). A study of spatial features of clones in a population of bracken fern, Pteridium aquilinum (Dennstaedtiaceae). Am. J. Bot., 80(5), 537–544. doi:10.1002/j.1537-2197.1993.tb13837.x

Quinlan, A. R. & Hall, I. M. (2010). BEDTools: a flexible suite of utilities for comparing genomic features. Bioinformatics, 26(6), 841–842. doi:10.1093/bioinformatics/btq033

Reusch, T. B. H., Boström, C., Stam, W. T. & Olsen, J. L. (1999). An ancient eelgrass clone in the Baltic. Mar. Ecol. Prog. Ser., 183, 301–304. doi:10.3354/meps183301

Reusch, T. B. H. & Boström, C. (2011). Widespread genetic mosaicism in the marine angiosperm Zostera marina is correlated with clonal reproduction. Evol. Ecol., 25(4), 899–913. doi:10.1007/s10682-010-9436-8

Reusch, T. B. H., Baums, I. B. & Werner, B. (2021). Evolution via somatic genetic variation in modular species. Trends Ecol. Evol., 36(12), 1083–1092. doi:10.1016/j.tree.2021.08.011

Schön, I., Martens, K. & van Dijk, P. (2009). Lost sex: The Evolutionary Biology of Parthenogenesis. Dordrecht: Springer Dordrecht.

Smith, M. L., Bruhn, J. N. & Anderson, J. B. (1992). The fungus Armillaria bulbosa is among the largest and oldest living organisms. Nature, 356(6368), 428–431. doi:10.1038/356428a0

Stoeckel, S., Arnaud-Haond, S. & Krueger-Hadfield, S. A. (2021a). The combined effect of haplodiplonty and partial clonality on genotypic and genetic diversity in a finite mutating population. J. Hered., 112(1), 78–91. doi:10.1093/jhered/esaa062

Stoeckel, S., Porro, B. & Arnaud-Haond, S. (2021b). The discernible and hidden effects of clonality on the genotypic and genetic states of populations: improving our estimation of clonal rates. Mol. Ecol. Resour., 21(4), 1068–1084. doi:10.1111/1755-0998.13316

The 1000 Genomes Project Consortium. (2015). A global reference for human genetic variation. Nature, 526(7571), 68–74. doi:10.1038/nature15393

The 1001 Genomes Consortium. (2016). 1,135 genomes reveal the global pattern of polymorphism in Arabidopsis thaliana. Cell, 166(2), 481–491. doi:10.1016/j.cell.2016.05.063

Van der Auwera, G. A. & O’Connor, B. D. (2020). Genomics in the cloud: using Docker, GATK, and WDL in Terra: O’Reilly Media.

Yu, L., Boström, C., Franzenburg, S., Bayer, T., Dagan, T. & Reusch, T. B. H. (2020). Somatic genetic drift and multilevel selection in a clonal seagrass. Nat. Ecol. Evol., 4(7), 952–962. doi:10.1038/s41559-020-1196-4

